# Phenotypic plasticity, life cycles, and the evolutionary transition to multicellularity

**DOI:** 10.1101/2021.09.29.462355

**Authors:** Si Tang, Yuriy Pichugin, Katrin Hammerschmidt

## Abstract

**SUMMARY:** Understanding the evolutionary transition to multicellularity is a key problem in evolutionary biology (1–4). While around 25 independent instances of the evolution of multicellular existence are known across the tree of life (5), the ecological conditions that drive such transformations are not well understood. The first known transition to multicellularity occurred approximately 2.5 billion years ago in cyanobacteria (5–7), and today’s cyanobacteria are characterized by an enormous morphological diversity, based upon which they have been classified into five sections. They range from single-celled species (section I), unicellular cyanobacteria with packet-like phenotypes, e.g., tetrads (section II) and simple filamentous species (section III) to highly differentiated filamentous ones (sections IV and V) (8–10). The unicellular cyanobacterium *Cyanothece* sp. ATCC 51142, an isolate from the intertidal zone of the U.S. Gulf Coast (11), has been classified as a section I species, and it phylogenetically clusters with the other N2-fixing unicellular cyanobacteria (12).

Here we report a facultative multicellular life cycle for a unicellular cyanobacterium, where multicellular filaments and unicellular stages alternate. In a series of experiments we identify the environmental factors underlying the phenotypic switch between the two morphologies. Then we experimentally confirm that the dissolution of filaments into solitary cells is triggered by changes in the external environment, which in turn is modified by the *Cyanothece* cells. Finally, using numerical models, we test a number of hypotheses regarding the nature of the environmental cues and the physical mechanisms underlying filament dissolution. While results predict that the observed response can be caused by an excreted compound in the medium, we cannot fully exclude changes in nutrient availability (as in (13,14)). The best-fit modeling results demonstrate a nonlinear effect of the compound, which is characteristic for density-dependent sensing systems (15,16). Further, filament fragmentation is predicted to occur by means of connection cleavage rather than by cell death of every alternate cell, which is corroborated by results from fluorescent and scanning electron microscopy. The phenotypic switch between the single-celled and multicellular morphology constitutes an environmentally dependent life cycle, which likely represents an important step en route to permanent multicellularity.

## RESULTS

### The filamentous morphology depends on the environment

To investigate the environmental factors that favor a multicellular morphology, we exposed the single-celled cyanobacterium *Cyanothece* sp. ATCC 51142 (hereafter *Cyanothece* sp.; Figure 1A) to the range of salinities and population densities it would encounter at its isolation site, the intertidal zone of the U.S. Gulf Coast (11). When culturing replicate populations of *Cyanothece* sp. (5*10^5^ cells/mL) in media with different amounts of added NaCl (0 - 300 mM), we observed that the occurrence of a filamentous morphology after 48 h (Figure 1A) significantly depends on the salinity of the medium (ANOVA, F_10,32_ = 953.10, *P*<0.0001). More specifically, at 0 mM NaCl, the whole population displayed a filamentous morphology, ranging from 4-celled up to 16-celled filaments, whereas at 300 mM NaCl (30 PSU), the whole population was single-celled. At intermediate salinities, we always observed both types - 4-celled filaments and single cells - with a higher fraction of filaments up to 90 mM NaCl, whereas at higher salinities single cells represented the most dominant morphology. Notably, we found that populations with a higher fraction of filaments also contain higher cell numbers/mL (ANOVA, F_1,32_ = 11.658, *P*=0.0018).

**Figure 1.**
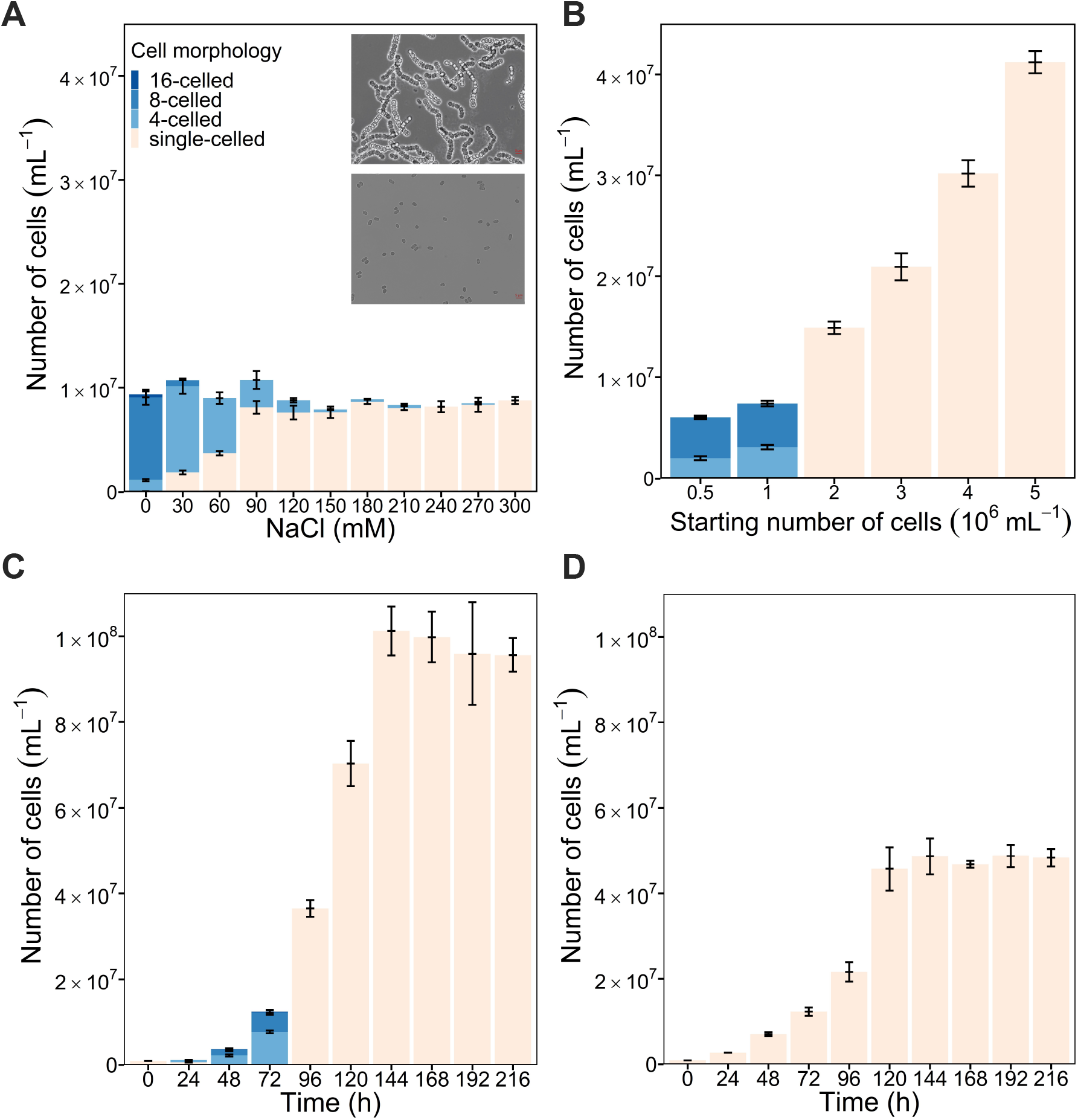
**Filamentous morphology of *Cyanothece sp*. ATCC 51142 depends on environmental salinity (at 48 h) (A), population density (at 48 h; in BG11 without added NaCl) (B), varies over time in batch culture (in BG11 without added NaCl) (C), and is not observed over time in BG11 with added NaCl (300 mM) (D). All experiments were performed in 10 mL media. Error bars represent standard deviations (of each sub-bar for A-C) (n=3).**

Next we addressed the role of population density for filament formation (in BG11 media without added NaCl). Replicate populations were initiated with six different starting population densities (all stemming from the same culture), ranging from 5*10^5^ – 5*10^6^ cells/mL. We found that starting cell density was a good predictor of *Cyanothece* sp.’s morphology – after 48 h we only detected filaments in cultures initiated with low density, whereas when initiated with higher starting cell densities (≥2*10^6^ cells/mL), we did not observe filaments (Figure 1B, X^2^ = 16.89, d.f. = 5, *P* = 0.0047).

To evaluate the effect of population density on population composition in more detail, we followed replicate populations inoculated with single-celled *Cyanothece* sp. of the low-density populations (5*10^5^ cells/mL) in 0 mM NaCl media for 5 days (Figure 1C). Already, after 24 h, we detected first 4-celled filaments. Thereafter, the whole population became filamentous, composed mostly of 4-celled and 8-celled filaments at 48 h, and of 4-celled, 8-celled, and 16-celled filaments 72 h post inoculation. Subsequently, the elongation of the filaments stopped, and from 96 h post inoculation onwards, we only observed single cells. We found that populations displaying the transient filamentous morphology grew faster and reached higher cell densities as compared to populations that only occur in the single-celled stage throughout their growth cycle (Figure S1; Figure 1D; ANOVA, F_1,5_ = 331.38, *P*<0.0001). Note that the population composition differed depending on the volume of the media; the previously described replicate populations were cultured in 10 mL each. While the same pattern and change between single cells and filaments were observed in volumes of 1 mL each, filaments were present for shorter times so that populations contained single cells already at 72 h post inoculation (Figure S2).

### Cyanobacterial morphology switches are mediated by external cues

The results so far indicate that the observed switches between single cells and filaments and back are density dependent - at higher cell densities, filamentation is inhibited and filament fragmentation is induced. This might be mediated by mechanisms of direct cell-cell contact or through depleted or excreted compounds in the medium. To test this, we harvested both supernatants from cultures inoculated with higher starting cell densities (5*10^6^ cells/mL) that did not produce filaments (filament inhibitor; harvested at 24 h after inoculation, Figure 1Figure 1B), and supernatants from cultures inoculated with low cell densities (5*10^5^ cells/mL) directly after filament fragmentation (filament fragmentor; harvested at 96 h, Figure 1C). To test whether compound (i.e. nutrient) depletion hinders filamentation, we added fresh culture medium (BG11) to both supernatants and to ddH_2_O, creating BG11 dilutions from 0 – 100 % with 20 % increments. Whereas we observed filaments at low media concentrations in ddH_2_O (20 % BG11, Table S1), we needed to add more BG11 to the two supernatants (60 %, 80 % BG11) to observe the multicellular morphology. As the lowest nutrient levels are found in the 20 % BG11:ddH_2_O mixture, where filament formation was observed (in contrast to the 60 % and 80 % BG11:supernatant mix), an overall depletion of nutrients as the factor hindering filamentation can be excluded. This notion is supported by the growth trajectory of single cells after fragmentation (Figure 1C), which show that cells can still reproduce (in fact they display the shortest generation time in the period directly after fragmentation, see Figure S1) and reach a high final concentration in the media post-fragmentation, and so strongly indicates that the post-fragmentation media (i.e. filament fragmentor) contains sufficient nutrients for growth. This can also be seen in the results from the density-dependent experiment (Figure 1B), which show that when starting with 5*10^6^ cells/mL, the cell density reached ∼ 4.1*10^7^ cells/mL after 48 h. Note that while we demonstrate that post-fragmentation media contains sufficient nutrients for growth, we cannot fully exclude the depletion of a specific compound required for maintaining growth in the filamentous form.

The harvested supernatants were also added to cultures of low-density single cells and to 48 hours-old filaments and compared to a control, i.e. fresh culture medium (BG11 without the addition of NaCl). After 24 h of incubation, we observed that both single cells and filaments displayed a different behavior when exposed to the two types of supernatants as compared to the control (Figure 2, ANOVA, F_3,35_ = 12.25, *P*<0.0001). While in the control, single cells formed filaments and filaments did not fragment, both supernatants inhibited filament formation from single cells, and also led the filaments to fragment. This strongly suggests that the phenotypic change is not related to direct cell-cell contact, but that substances or the depletion of substances in the supernatant affect the transition between both phenotypes. Interestingly, both supernatants can be used interchangeably – both are able to inhibit filament formation from a single-celled ancestor and are also able to induce filament fragmentation. However, whether both supernatants contain or lack an identical substance needs to be investigated.

**Figure 2.**
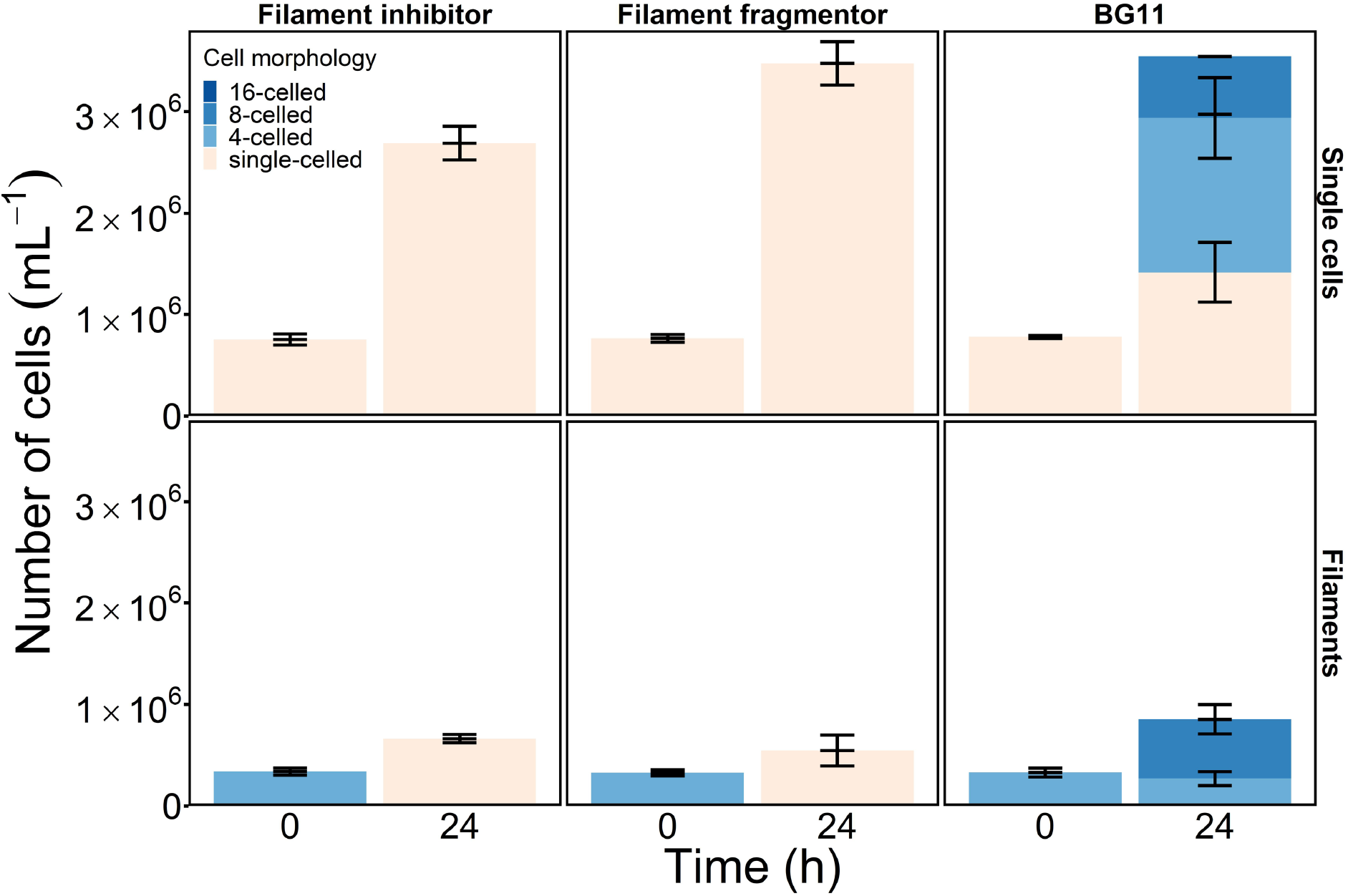
**Cell morphology depends on the external medium. Filament inhibitor and fragmentor supernatants stop the formation of filaments from single cells, and lead to the dissolution of existing filaments within 24 h of exposure. This is in contrast to the control, untreated freshwater medium (BG11), which induces filament formation from single cells and filament elongation. Error bars represent standard deviations of each sub-bar (n=3).**

As we only observed comparatively short filaments, i.e. with a maximum of 16 cells, before they disintegrated, we investigated whether this is due to culture density (mediated by substances or the lack of substances within the supernatant) or whether it is also modulated by other factors that might constrain filament length. To test this, we diluted cultures containing filaments at 72 h by adding fresh culture media. We found that in diluted cultures, filaments increase in lengths (Figure S3), indicating that the density of the culture affects filament length.

### Mechanisms of filament dissolution and compound action

We use the collected data to fit a series of models to compare different hypotheses regarding (i) the mechanism of filament dissolution, i.e. cell death or cleavage of the connection sites, and (ii) the dynamics of the compound, i.e. whether the observed pattern is caused by the accumulation or consumption of a compound. The theoretical modeling approach allows us to explore a much wider set of parameters in different combinations as compared to empirical work.

We consider three families of models. The first family, the *toxic compound models*, assumes that cells produce a compound inducing cell death leading to the fragmentation of filaments. The second family, the *disconnecting compound models*, assumes that cells produce a compound inducing cleavage of connections between cells. The third family is complementary to the previous one: these models also assume that filaments fragment due to connection loss, which is caused by consumption of an initially present compound that stabilizes connections. To highlight this complementarity, this family is referred to as *connecting compound models*. In each family, we consider a number of models, which differ by the character of the compound action (rate of cell death or connection loss) with respect to the compound concentration. For example, we compare individual models, where the action rate proportionally depends on the concentration, with step-dependence models, where the compound has no effect at concentrations below a certain threshold. In total, we considered 12 models each for the toxic and disconnecting compounds, and 8 models for the connecting compound (see Figures S4 and S5, Tables S2 and S3 for the complete list). These 32 models take into account a much more diverse set of possible dynamics of environment-dependent filament fragmentation than those used in theoretical studies so far (17), see also Supplementary Text S1.

We found that (dis-)connecting models in which fragmentation is caused by the loss of cell connections fit significantly better than the toxic compound models resulting in cell death (one-sided Mann-Whitney U test between regression errors of toxic and disconnecting compound models yields U = 739355, at n_toxic_=3000, n_disconnecting_=2884, with p < 10^−10^ and the same significance level was found when we contrasted toxic and connecting compound models), see also Figure 3A. Thus, we conclude that the observed population dynamics seem to be caused by cells disconnecting from each other rather than an outcome of cell death compromising filament integrity. At the qualitative level, the poor fit of toxic compound models comes from the inability of these models to sustain a purely unicellular population – there must be multicellular filaments present in any stable population under the influence of a toxic compound. Specifically, we show that if filament fragmentation is caused by cell death, then half of the population must be filamentous (see Supplementary Text S2) – something we do not observe in the experiment. Furthermore, even if we ignore the distribution of filaments sizes, and assume that the compound selectively kills every second cell in each filament, such a fast transition from the filamentous population to the one made of solitary cells, as we observed, means that the population must suffer a massive loss. Yet, we don’t observe any significant change in the population growth rate at the moment of filament dissolution (see Figure S8).

**Figure 3.**
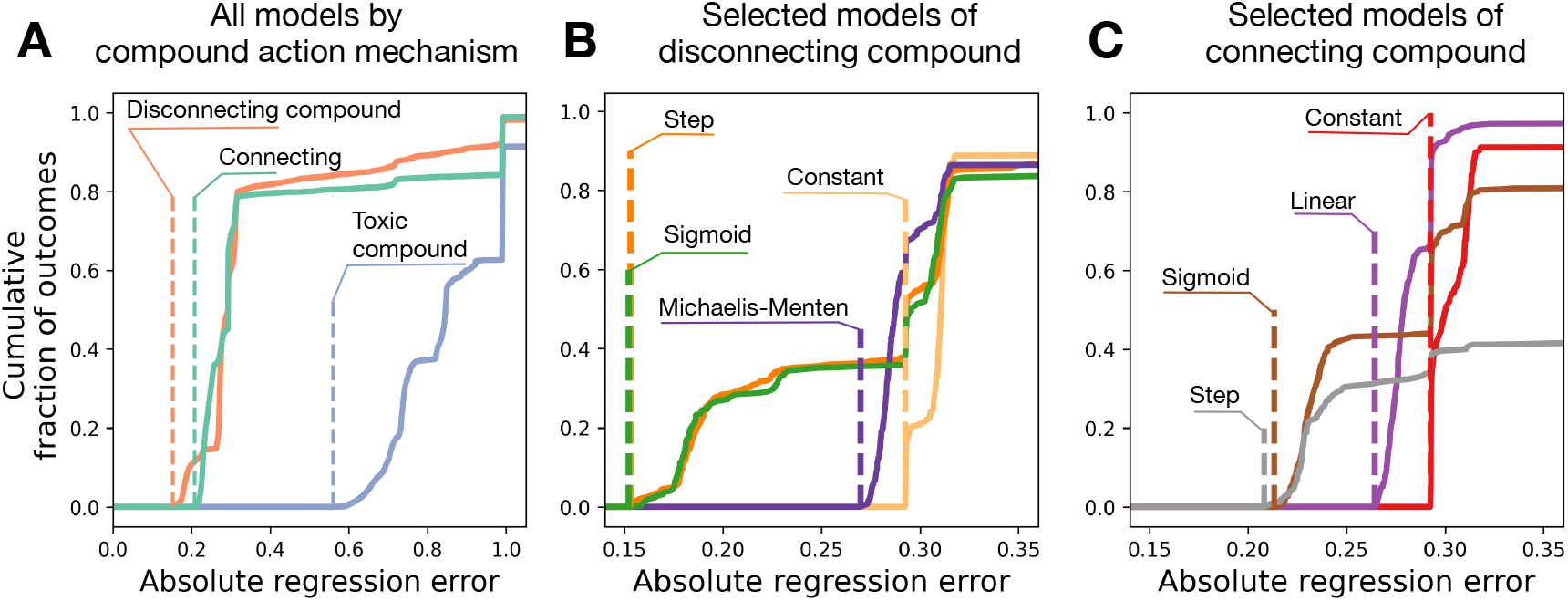
The mechanism of filament dissolution is likely the cleavage of cell connections (A) with a strong non-linear response to active substance concentrations (B, C), as suggested by modeling. Plots show sample cumulative distribution functions of regression errors from 250 independent optimizations for each model. Dashed lines represent the minimal regression error in each group. **(A)** Models in which connections are destroyed due to production (orange, 12 models in total) or consumption (green, 8 models) of a (dis)connecting compound provide much smaller regression errors than models with a toxic compound (blue, 12 models). **(B)** For models with a disconnecting compound (the best class overall), the smallest regression errors were observed among models in which the rate of connection cleavage is negligible at compound concentrations below a certain threshold (e.g. in the step and sigmoid models). **(C)** For models with a connecting compound, the smallest regression errors were observed among models in which the rate of connection cleavage can quickly rise with the compound depletion (e.g. also in the step and sigmoid models). Only four models from each class are shown, for the complete set of models see Figure S6 and Figure S7.

This finding is corroborated by empirical evidence, where we did not observe cell death in the experimental cyanobacterial populations (dead cyanobacterial cells can be identified by the lack of a fluorescent signal when populations are observed using fluorescent microscopy), which provided the data for the models reported here. In addition, using scanning electron microscopy (SEM) on 72-hour-old filaments, we found that the size of the connecting areas between cells varied greatly. While some cells share a larger area with their neighboring cell, other connections are thinner and seem to disconnect easily (see example filament with breaking connection in the center of Figure S9). We suspect that the closely connected cells are cells that are undergoing (or have just completed) cell division, and that the cells with weaker connections underwent cell division in previous rounds. Presumably, during filament fragmentation, filaments dissolve due to the breaking of these weak connections.

Comparing the remaining models (i.e., connecting and disconnecting families), we found that the most accurate fit is achieved by these disconnecting compound models, where the effect of the compound action is close to zero when the concentration of the compound is below a threshold: in step and sigmoid models (Figure 3B) and also in fracture, and in breaking point models (Figures S4, S6). There is no significant difference between the regression errors achieved in these four models. The remaining models demonstrate much larger regression errors, similar to those of control models (constant and proportional), see Table S4.

Among the connecting compound models, three out of five best models explicitly feature a threshold (step, sigmoid, and quadratic concave), see (Figure 3C). Two more models with similarly good fit do not have an explicit threshold designed into them (inverse and exponent). Still, all five models share a rapid decrease in filament fragmentation rate at a low compound concentration, followed by a plateau of low sensitivity to concentration – a feature absent among poor fit models (constant, linear, quadratic convex), see Figures S5, S7 and Table S5.

Altogether, connecting and disconnecting models show that in order to observe patterns found in the experiments, the rate of filament fragmentation must depend non-linearly on the compound concentration. In all best-fitted models, the fragmentation rate skyrockets at some moment: either after exceeding a threshold compound concentration for the disconnecting compound models, or after expiration of the consumed compound for the connecting models.

## DISCUSSION

### Phenotypic heterogeneity in *Cyanothece* sp. ATCC 51142

When exposing the unicellular cyanobacterium *Cyanothece* sp. ATCC 51142 to a range of salinities comparable to conditions it would experience in its native habitat, the intertidal zone, we observe two distinct microbial lifestyles: the previously described single cell stage (11), and a newly observed filamentous stage (Figure 1A). At low population densities, the single cell stage predominates in cultures with higher salinities (≥90 mM NaCl), while the filamentous stage is most prevalent under low-salinity conditions and the only stage in the populations cultured in the salinity-free environment. The repeatability and the high rate of the filamentous stage in the population exclude genetic changes, such as mutation or gene amplification (18), as the underlying mechanisms for the change in morphology. Moreover, the co-existence of both morphological stages at the intermediate salinities is indicative of phenotypic heterogeneity.

Phenotypic heterogeneity has been reported for many bacteria and is defined as diverse phenotypes arising from genetically identical microbes that reside in the same microenvironment (19). Several molecular mechanisms underpin phenotypic heterogeneity, such as stochastic state switching, periodic oscillations, cellular age, and cell-to-cell interactions. Whereas in this study, stochastic state switching can be excluded due to the exclusiveness of either of both stages in the two extreme environments (0 mM and 300 mM NaCl), periodic oscillations and cellular age can be excluded when the populations in the different salinities are compared, as all populations were initiated from the same culture. This promotes cell-to-cell interactions as the most likely mechanism underpinning the switch between the two distinct phenotypes. This is confirmed by the outcomes from the experiment, where the effect of starting population density on lifestyle was investigated. Here, the filamentous stage was only observed at lower population densities (≤1×10^6^ mL^-1^), whereas at higher densities only single cells were detected. This indicates that an increase in cell-to-cell interactions prevents the formation of the filamentous stage. Cell-to-cell interactions could either happen via direct contact between cells (20), or through indirect means, for example in response to cues from other cells that are excreted into the medium, e.g. quorum sensing (21), or through the consumption of nutrients. This was addressed by adding cell-free supernatants from high-density to low-density cultures, which indeed inhibited filament formation, thus confirming that filament inhibition is not caused by direct cell-cell interactions.

### Alternating phenotypes constitute an environmentally mediated life cycle

While the different microbial lifestyles and transitions between them have been researched in much detail, this approach only provides a fragmented picture of microbial life cycles that occur in nature (22). When we follow low-density populations of *Cyanothece* sp. in the no-salinity environment over time, a microbial life cycle with alternating single cell and multicellular life stages, can be observed as described for other single-celled bacteria, such as *Bacillus subtilis* that transition through single cell, filamentous, and dormant life stages (22) or for experimental populations of *Pseudomonas fluorescens* (23). In the case of *Cyanothece* sp., 24 h after initiation of the single-celled culture, filaments can be observed that constitute the only cell-stage for another 48 h. Thereafter, within the next 24 h, filaments disappear so that single cells are the only stage present in the population at 96 h. The disintegration of filaments is central to the completion of the life cycle. We have experimentally shown that filament dissolution is triggered by changes in the environment caused by the *Cyanothece* population. For example, at high initial density of solitary cells, the environment is modified so quickly that the filaments do not get enough time to form at all. In addition, when the supernatant from such a highly dense population is added to a population composed of filaments, they dissolve (see Fig. 2).

To learn more about the nature of the compound and about the way filaments transitioned to single cells, i.e. whether every alternate cell dies or whether cells separate through cleavage of connection sites, the experimental data was fitted with 32 different models that vary in the effect of the compound on the filament (cell death or connections cleavage), and in the effect of the concentration of the cue. As the models that assumed cell death resulted in a worse fitting than the ones where the cue resulted in the cleavage of connection sites, filaments likely disintegrate into single cells without cell death.

The classic example for phenotypic switching induced by cues in the medium is quorum sensing (21). The known mechanisms for quorum sensing involve cell-to-cell signalling by molecules called autoinducers. They are involved in a regulation pathway featuring a positive feedback loop – the more autoinducer molecules are present, the higher is their production rate (15,16). As a consequence, the kinetic models of quorum sensing feature bi-stable dynamics: the cell is either on or off. While the precise kinetics and regulation of the compound lie outside of the scope of our work in the context of this study, the known kinetics models of quorum sensing is best represented by the step model (Figures S4 and S5). The sigmoid model is the generalization of the step model and features it as a limiting case. These two models demonstrate the two best fits in both model families: connecting and disconnecting compound (Tables S4 and S5).

From a theoretical perspective, multicellular life cycles with fission into multiple pieces have a selective advantage when a fragmentation event is costly in some way (24,25). A typical mechanism making fragmentation costly is cell death observed in some species of filamentous cyanobacteria, where specific cells, necridia, undergo programmed cell death, when releasing hormogonia (motile reproductive filaments) from the mother filament (26). The modeling results in combination with results from fluorescent and scanning electron microscopy suggest however that cell death does not happen here. The next alternative is that either production of inhibitor/fragmentor or the response to its high concentration is costly. Given that the results of the fitting indicate a more complex response of *Cyanothece* sp. to inhibitor/fragmentor than the simple mass action law, this supports the hypothesis that some dedicated mechanism, such as quorum sensing, is involved in the observed dissolution of filaments. Within cyanobacteria, our observation seems to be most similar to cyanobacteria that have traditionally been classified as belonging to section II (unicellular and organized into packet-like structures, (10)). It has been long-known that these species rarely occur as solitary cells, but mostly as micro-or macroscopic, irregular, formless, or granular colonies (27), but there has been a recent report that one species belonging to this group also displayed morphological phenotypic plasticity – transitioning between single cells, dyads (2-celled colonies), triads (3-celled colonies), tetrads (4-celled colonies) and colonies with more than four cells - depending on the chemical composition of the culture medium and depending on their growth stage (28). While in our case we also observe morphological phenotypic plasticity depending on the environment, we see the formation of filaments (the underlying morphology within the cyanobacterial sections with complex multicellularity) for a unicellular (section I) cyanobacterium, which has been classically described to occur as single cells (10,11).

### Internalization of the regulation of phenotypic plasticity may drive the evolutionary transition to multicellularity

The environmentally dependent filamentation/fragmentation of *Cyanothece* sp. allows drawing parallels between the population dynamics observed in our study and a life cycle of a multicellular organism. The unicellular phenotype observed in the saline environment is analogous to the start of a life cycle from a single cell/propagule. The process of filamentation in low-salinity conditions represents the growth of a multicellular organism. Finally, the fragmentation of filaments into single cells is comparable to multicellular reproduction resulting from new propagules being released into nature.

While there is no evidence that the repeated filamentation and fragmentation occur under natural conditions, we can speculate that the observed effect might be at the core of an environmentally dependent life cycle. Such a hypothetical life cycle would start once single cells of the marine *Cyanothece* sp. find themselves in a compartment with reduced salinity, for example a river estuary. There, through filamentation multicellular chains would be formed until high local densities are reached. Overcrowding would result in the fragmentation of filaments into independent cells. Given the small size of single cells compared to multicellular filaments, the newly released single cells are much more likely to be moved away from the overcrowded environment – either into the sea or to another freshwater compartment, ready to restart the cycle again.

While this is the first report of such an environmentally dependent life cycle for unicellular cyanobacteria, such life cycles are not only known for bacteria, such as *Bacillus subtilis*, which also alternates between motile single cell and filamentous stages (22). The slime mold *Dictyostelium discoideum*, for example, exists as single cells under favourable conditions, which aggregate into a multicellular slug capable of locomotion upon nutrient depletion (29). Eventually, the slug differentiates into stalk and fruiting body, releasing spores from which new single cells hatch. Another example is the predominantly unicellular marine choanoflagellate *Salpingoeca rosetta*, which forms multicellular colonies in the presence of prey bacteria (30). In all cases, the transition from the unicellular to a multicellular stage is a response to environmental cues. Moreover, even stages of highly sophisticated and integrated developmental animal life cycles have been discovered to depend on external factors and ecological triggers, some of which are based on communication with external bacteria (31). Theoretical models of life cycle evolution have shown that a changing environment can lead to the evolution of complex life cycles in which some cells live and reproduce as unicellular beings, while others form groups (32).

The transition between the different stages of a life cycle might initially be dependent upon the environment and be based on phenotypic plasticity, but recent findings suggest that the integration of these stages, i.e. the endogenization of life cycles, is central to the evolutionary transition to multicellularity (33,34). One could imagine an ancestral plastic response present in the unicellular ancestor that is co-opted, such as the density-dependent switch between single cells and filaments reported here. In that case, the transition from the predominantly unicellular life cycle with facultative multicellular stages to an obligate multicellular life cycle might be straightforward. Given an adequate selective environment, it would simply involve a change from a facultative to an obligate expression of the underlying genes, for example of the ones that are involved in filament formation. This might result in filaments that do not completely disintegrate but where, e.g., only cells at the end(s) of the filament get released. As a result, both tasks, i.e. growth of the filament and release of propagules, would happen simultaneously - one would observe a differentiation in the form of reproductive division of labour.

Even though, this life cycle has been artificially induced in the laboratory, it seems to be realistic for what is happening in nature. Many habitats are characterised by rapidly changing environmental conditions, so for example, a unicellular organism isolated in one environment might possess a completely different phenotype/life stage in another. While it is tempting to classify organisms based on their phenotypes, it is important to realise that in the laboratory we often investigate only parts of an organism’s life cycle. For studying the transition to multicellularity, the importance of the morphological and physiological flexibility of the unicellular ancestor is becoming more and more apparent, for example when studying the complex life cycles of protozoans to infer the last unicellular and last common multicellular ancestor of animals (31) or when comparing experimentally evolved nascent stages of early multicellular life cycles to the highly differentiated life cycles and cells of their closest multicellular relatives. Thus, at least some (if not all) transitions to differentiated multicellularity might have been a rewiring from temporal differentiation of life cycle stages to spatial differentiation in multicellular organisms (35). This poses important questions regarding the ease of such transitions, the ease of reversals to unicellularity, but also regarding our views of the transition to multicellularity (36). Should we really think about the transition to clonal multicellularity as the multi-step process that starts with the evolution of undifferentiated multicellularity getting more complex over evolutionary timescales or rather as a one-step process starting with a complex unicellular ancestor that directly transitions into a differentiated multicellular organism?

## SUPPLEMENTAL INFORMATION

The simulation code, its results, and data processing are publicly available at https://github.com/yuriypichugin/cyanobacteria-filament-fragmentation.

## ACKNOWLEDGEMENTS

We thank Peter Deines, Caroline Rose and Nancy Weiland-Bräuer for critical comments on the manuscript. ST was funded by the China Scholarship Council (CSC), Beijing, China. YP is grateful to the Max Planck Society for generous funding. K.H. thanks the Hamburg Institute for Advanced Study (HIAS) and the Joachim Herz Foundation for support.

## AUTHOR CONTRIBUTIONS

ST and KH designed the experiments. ST performed the experiments. YP performed the theoretical modeling. All authors interpreted the results and wrote the manuscript.

## METHOD DETAILS

### Bacterial strain and culture conditions

*Cyanothece* sp. ATCC 51142 (recently reclassified as *Crocosphaera subtropica* (37)) was obtained from the ATCC culture collection. Cells were grown photoautotrophically at continuous light with a light intensity of 30 μmol m^-2^ s^-1^ in liquid culture at 30°C. To evaluate the effect of salinity on the morphology of *Cyanothece* sp., we carried out a growth experiment, where we supplemented BG11 media with NaCl. In total, we tested three replicates in each of the eleven NaCl concentrations (0, 30, 60, 90, 120, 150, 180, 210, 240, 270, 300 mM) with a total volume of 10 mL. We recorded the population composition 48 h after the start of the experiment. To test for the effect of starting population density on the morphology of *Cyanothece* sp., we created a gradual series of starting population densities (5*10^5^, 10^6^, 2*10^6^, 3*10^6^, 4*10^6^, 5*10^6^ cells/mL, a total volume of 10 mL), with three replicates per condition. We again recorded the population composition 48 h after the start of the experiment.

### Supernatant test

To test whether the phenotypic switch is mediated by nutrient depletion, fresh culture medium (BG11) was diluted with filament inhibitor, filament fragmentor and ddH_2_O, creating a gradual series of BG11 ratios (0 %, 20 %, 40 %, 60 %, 80 %, 100 %), within a total volume of 1 mL. Low-density single cells (5*10^5^ cells/mL) were cultured in all combinations in the 24-well plate, with three replicates for each combination. After 48 h, the morphology of the cells within each replicate was observed under the microscope and the presence of filaments was recorded.

To investigate whether filament fragmentation depends on cellular age, single cells were grown in 10 mL BG11 in tissue culture flasks for 72 h, after which filament formation was confirmed under the microscope. Cultures were gently mixed, and 20 μL of the filamentous *Cyanothece* sp. population was transferred to 980 μL of fresh BG11 medium. As a control, 1 mL of culture without the addition of fresh medium was grown in parallel. Each treatment was carried out with three replicates. After 24 h, replicate populations (n=3 each) were quantified. To differentiate whether the morphology changes from single cells to filaments and back have been induced by direct cell-cell contact or through (excreted) compounds in the media, we harvested two supernatants: (i) filament inhibitor, and (ii) filament fragmentor. We created the filament inhibitor supernatant by setting up replicate cultures (n=3) with high starting cell densities (5*10^6^ cells/mL) in BG11 media. We let them grow for 24 h under the conditions described above, after which we centrifuged samples (at 20 × g for 3 minutes). Thereafter we processed the supernatant through a 0.22 μm filter (Syringe filter, membrane: PES) to exclude cyanobacterial cells. The process for harvesting the filament fragmentor supernatant was the same, only that the culture was initiated differently. Here, low-density cultures (5*10^5^ cells/mL) (n=3) were set up in BG11 and closely monitored. We harvested the supernatant directly after filament fragmentation occurred. Directly after harvesting the supernatants, we set up an experiment, where we exposed replicate populations (n=3 each) of single cells (5*10^5^ cells/mL) and 48-hours-old filaments to either filament inhibitor or filament fragmentor supernatants or to fresh BG11 media. To achieve this, 1 mL unicellular culture (exponential phase) or 1 mL of 48-hours-old filaments were centrifuged, and the resultant supernatants were discarded. Thereafter cells were resuspended in 1 mL of either filament fragmentor supernatant, filament inhibitor supernatant, or fresh BG11, respectively. We recorded the population composition 24 h after the start of the experiment.

### Quantification of population density and composition

Population density and composition was quantified with a cell counting chamber (Neubauer improved, depth: 0.1 mm) from which digital photographs were taken (camera AxioCam MRR3 mounted to the microscope ZEISS Imager.M2m). More specifically, from each replicate population (total volume of 1mL) 1 μL was assayed, and cells within the final volume were calculated with the formula: Cells in 1μl = (number of cells in the main square) (1μl) /0.004. Images were counted manually using the software ImageJ, and classified into four categories: single cells, 4-celled filaments, 8-celled filaments and 16-celled filaments. In addition, we screened the populations for cell death using fluorescent microscopy (dead cyanobacterial cells can be identified by the lack of a fluorescent signal).

### Scanning electron microscopy (SEM)

Single cells were grown in BG11 for 72 h, after which filaments were harvested by centrifugation, washed twice and resuspended in sterile PBS. Then samples were lyophilized for 12 h prior to imaging. The dried filaments were sputter-coated with a gold/palladium thin film for 100 s and a 15 mA current, using a Q150R sputter coater (Quorumtech, UK). Samples were imaged with a field-emission SEM and 10 kV electron beam (Zeiss Sigma 300, Zeiss, Germany).

### Statistical analysis

Sample size was chosen to maximise statistical power and ensure sufficient replication. Assumptions of the tests, that is, normality and equal distribution of variances, were visually evaluated. Non-significant interactions were removed from the models. All tests were two-tailed. Effects were considered significant at the level of P < 0.05. All statistical analyses were performed with JMP 9. Graphs were produced with R Studio Version 1.4.1564 and Python Matplotlib library.

### Theoretical model

In the model, we consider a population composed of filaments of different length. After division, cells always stay together, increasing the length of the filament. Cell divisions in each filament occur synchronously, so the filament length doubles at each division event. However, filaments divide independently from each other, hence division events among different filaments are not synchronized. The rate of cell division in density dependent -- the more cells are present in the population, the slower is cell division:

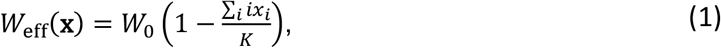

where *W*_eff_(**x**) is the division rate of cells in the population **x**, *W*_0_ is the maximal division rate, *x*_*i*_ is the number of filaments of the length *i*, and hence, ∑_*i*_ *ix*_*i*_ is the total number of cells in the population, *K* is the maximal number of cells that can be sustained in the population (carrying capacity). The maximal possible length of the filament in the model was limited to 32 cells, which is larger than any empirically observed filament. Filaments that reach that maximal size stop dividing.

We assume that filaments fragment due to the changes in the environment caused by the presence of cells. Here we consider three families of models. In **toxic compound** models cells produce a compound causing cell death. In **disconnecting compound** models cells produce a compound causing connections cleavage. In **connecting compound** models cells consume a compound that underlies filamentation. In both disconnecting and connecting models, the fragmentation occurs via loss of cell connections but in the first case cells produce the mediating compound, while in the second case they consume it.

In the disconnecting and toxic compound models, cells produce a compound *T*, which causes filament fragmentation. Each cell produces the compound with the unit rate. Produced compound decays with the rate *D*_comp_. Hence, the compound dynamics is given by

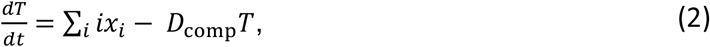

where the first term describes the production of compound by cells, and the second term describes the compound decay. The rate of compound production is set to one without loss of generality, as it is just a scaling factor for non-observed compound concentration.

In the connecting compound models, a fresh media initially contains a unit concentration of a compound, while cells consume the compound. Hence, the compound dynamics are governed by a different law

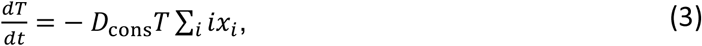

where *D*_cons_ is the rate at which a single cell consumes the compound.

In all cases, population dynamics are described by the set of differential equations

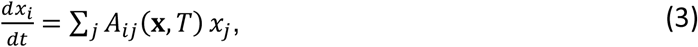

where the projection matrix *A*_*ij*_(**x**, *T*) shows the rate at which filaments of length *i* emerge from the filaments of length *j* by means of growth and fragmentation.

For the connecting and disconnecting compound models, filaments fragment by loosing connections between cells. Hence, the elements of this matrix are

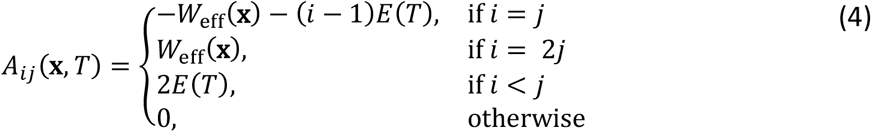

There, the first line describes the disappearance of filaments due to the growth or fragmentation, the second line describes the emergence of twice-longer filaments at each growth event, and the third line describes the emergence of shorter filaments after fragmentation. *E(T)* is the rate of effect of compound at the concentration *T*. In the disconnecting compound models, *E(T)* is a monotonically increasing function – more compound leads to higher rate of connections loss. In the connecting compound models, *E(T)* is a monotonically decreasing function – less compound leads to less stable connections.

For toxic compound models, filaments fragment whenever an internal cell dies. Hence, the elements of this matrix are

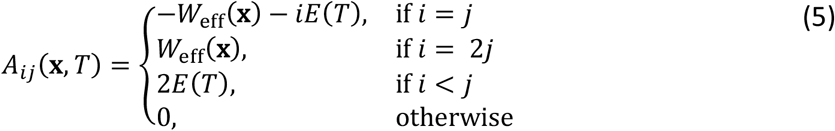

Note that the only difference between Eqs.(4) and (5) is the coefficient before *E(T)* in the first line. This represents that in a filament of *i* cells, there is *i-1* connections, which can be severed (if the fragmentation is due to the cleavage of connections) but *i* cells that can die (if the fragmentation is due to the death of cells).

For each family, we consider a number of models of the compound effect *E(T)*, see Figures S4 and S5 and Tables S2 and S3. There are 32 models in total. In each family, two control models represent situations, where the mechanism of the compound action is straightforward: the proportional model assumes a mass action law of the interaction between the compound and cells, the constant model assumes a spontaneous fragmentation of filaments, i.e. the compound plays no role. Other models represent situations, where the compound acts on filaments in a more complicated way.

### Data fitting and regression results

Four series of experiments are taken into account in the simulations. The first data set is the population composition at different starting densities (Figure 1B). The second data set is the population composition over time (Figure 1C). The third data set is the filament elongation test (Figure S3). The fourth data set is the investigation of the supernatant effects (Figure 2). For each tested combination of parameters, the simulations imitating experimental protocols were conducted.

To simulate the experiment shown in Figure 1B, the population was initialized with solitary cells at given concentrations and the population composition after 48 h recorded for comparison with experimental observations. To simulate the experiment shown on Figure 1C, the population was initialized with solitary cells at the given concentration and the population compositions at time points 24, 48, 72, 96, and 120 h after initialization were recorded. To simulate the experiment shown on Figure 2, first, the population was initialized with high density (filamentation inhibitor), or low density (fragmentation inducer) of solitary cells, and simulated for 24 or 72 h, respectively. Then, the concentration of the compound was sampled and used in the second simulation series, initialized with populations, given by records at 0 h in each of the sub-experiments. In each case, the composition after 24 h was recorded. To simulate the experiment shown in Figure S3, the population was initialized with the record given for 0 h and the population state after 48 h was recorded.

In each simulation, the mean square deviation between the experimentally observed and numerically simulated population composition was computed. Minimization of this value by adjusting growth and fragmentation parameters was the target of the fitting.

The experimental data has been fitted with each of 32 numerical models (see Tables S2 and S3 for definitions). The initial values of model parameters have been drawn randomly, so different runs of optimization ended at different points in the parameters space. To compensate for that, 250 independent optimization runs were computed for each model.

Regression errors were scaled by the error provided by the static population. This means that the fitting error provided by the population, which neither grows nor dies and is not affected by the compound, is equal to one. The hypothesis of a static scenario is clearly incorrect; therefore all fitting results with regression errors above one were discarded from the further analysis as completely unrealistic.

All toxic compound models demonstrated much larger regression errors than (dis-) connecting compound models (Figure 3A and Table S4). Among disconnecting models, four models demonstrated similar and low minimal regression errors (0.15 – 0.17): step, sigmoid, fracture, and breaking point. Quadratic model has shown the minimal error around 0.217. The remaining seven models (constant, proportional, linear, top-capped, bottom-capped, Michaelis-Menten, and saturating exponent) resulted in larger errors (0.26 – 0.29), see Figure S6 and Table S4. Among connecting compound models, the lowest regression errors (0.20 – 0.22) were observed for models capable to demonstrate a sudden increase in fragmentation rate at low compound concentrations (step, sigmoid, exponent, inverse, quadratic concave). At the same time, models in which the fragmentation rate increased gradually with compound loss (constant, linear, quadratic convex) resulted in larger regression errors (0.23 – 0.30), see Figure and S7, and Table S5.

## Supplemental Information

**Figure S1.**
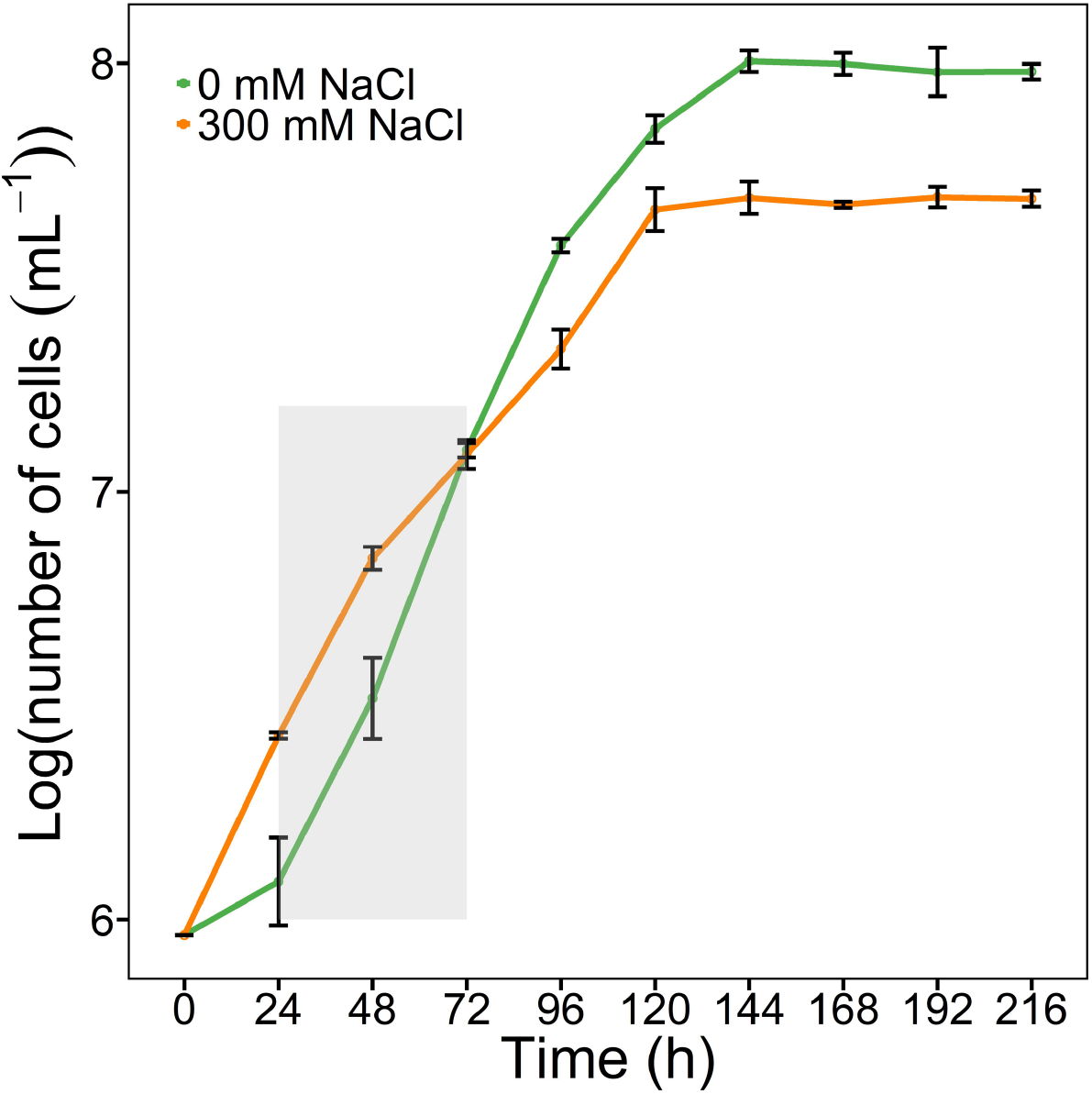
Comparison of the growth trajectory over time in batch culture in 10 mL BG11 medium with 0 mM and 300 mM NaCl. The grey area indicates the time period when filaments were observed in medium with 0 mM NaCl. The shortest generation time in freshwater is G_0 mM_ = 15.2 h (from 72 h to 96 h), while the shortest generation time in the highest salinity is G_300 mM_ = 17.5 h (from 24 h to 48 h).

**Figure S2.**
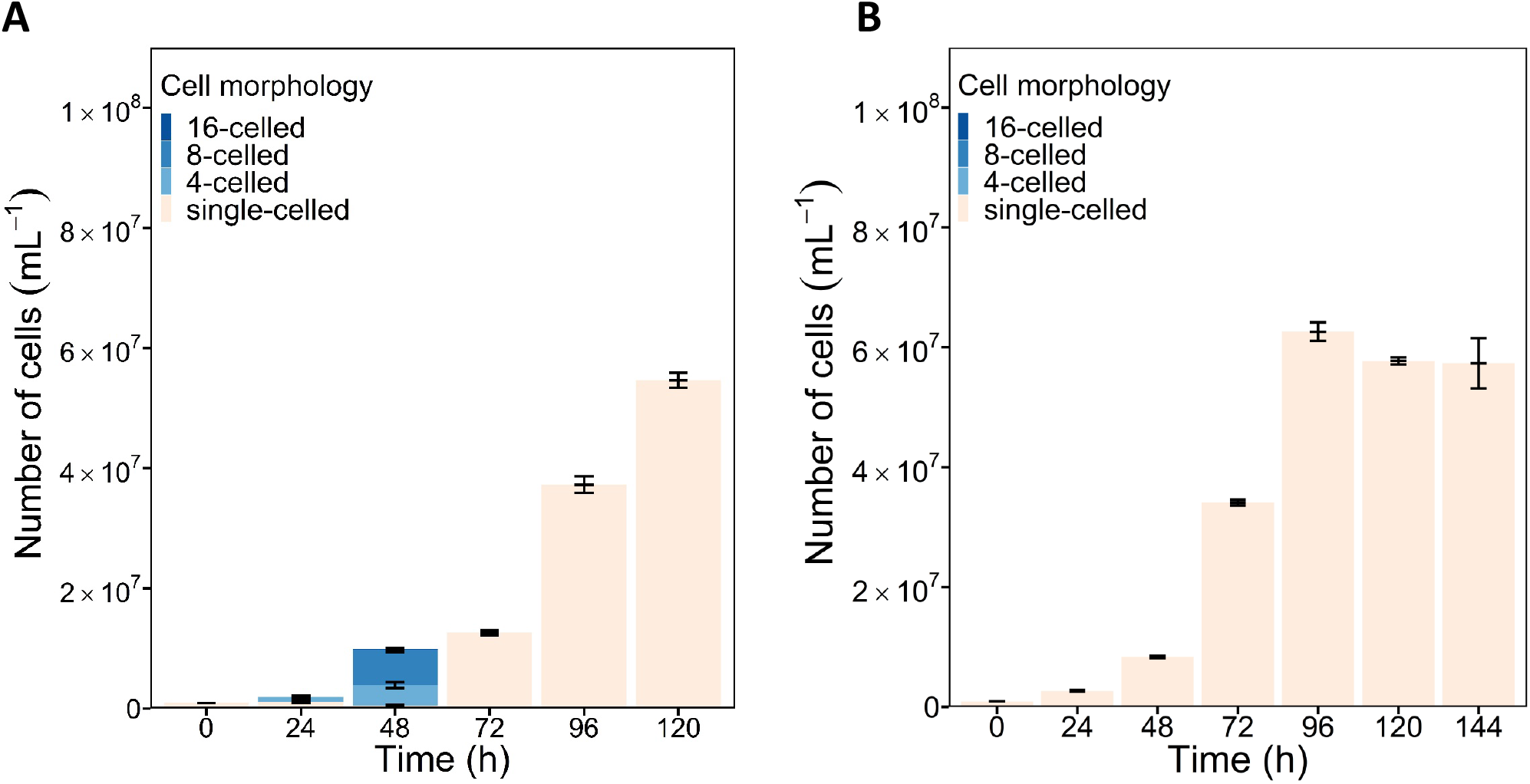
Population dynamics over time in 1 mL volume (24-well plates) in BG11 (A) and in BG11 with 300 mM added NaCl (B).

**Table S1.** The morphology of *Cyanothece* sp. is dependent on the composition of the medium. Fresh culture medium (BG11) was added to ddH_2_O and both supernatants, creating BG11 ratios from 0 – 100 % with 20 % increments. The emergence of the filamentous morphology was recorded after 48 h, starting with single cells of *Cyanothece* sp. in each dilution treatment. “+” represents filament occurrence; “-” represents no filament occurrence. While 20 % of BG11 in ddH_2_O provided sufficient nutrients for filament formation, 60-80 % of the BG11 was necessary to dilute the filament fragmentor/inhibitor medium before filaments were observed.

|  | BG11 ratio |  |  |  |  |  |
| --- | --- | --- | --- | --- | --- | --- |
|  | 100% | 80% | 60% | 40% | 20% | 0% |
| ddH<sub>2</sub>O | + | + | + | + | + | - |
| Filament fragmentor | + | + | + | - | - | - |
| Filament inhibitor | + | + | - | - | - | - |

**Figure S3.**
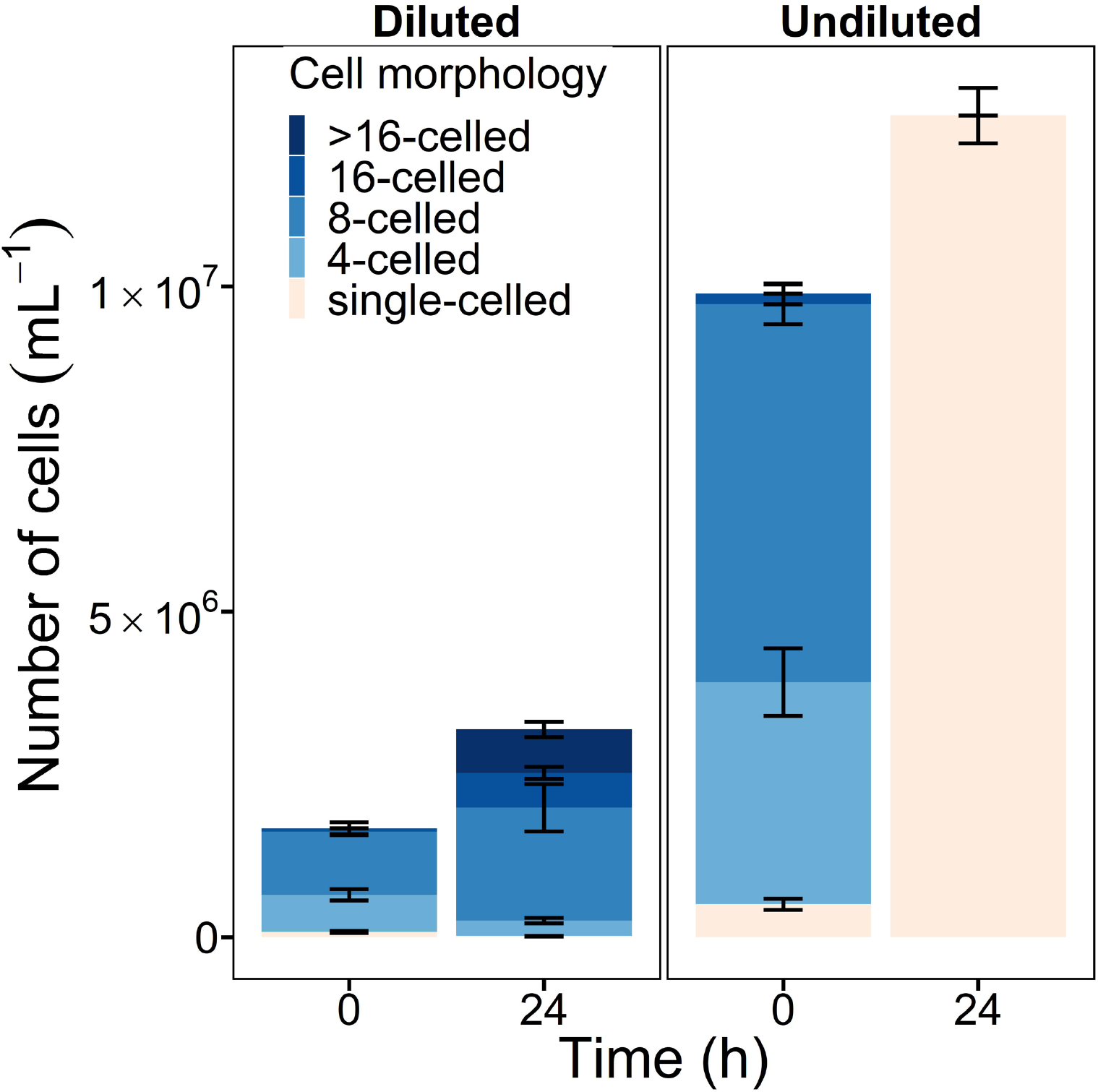
Population composition after the transfer of 72h-old filaments to new medium (left) in contrast to the original population (both in BG11 without added NaCl). When diluted, filaments kept growing and increased in length, indicated by the observation of filaments of longer than 16 cells in length and by a significantly higher proportion of 8-celled filaments, in contrast to the original culture, where 24 h later only single cells were observed. Error bars represent standard deviation of each sub-bar (n=3).

**Figure S4.**
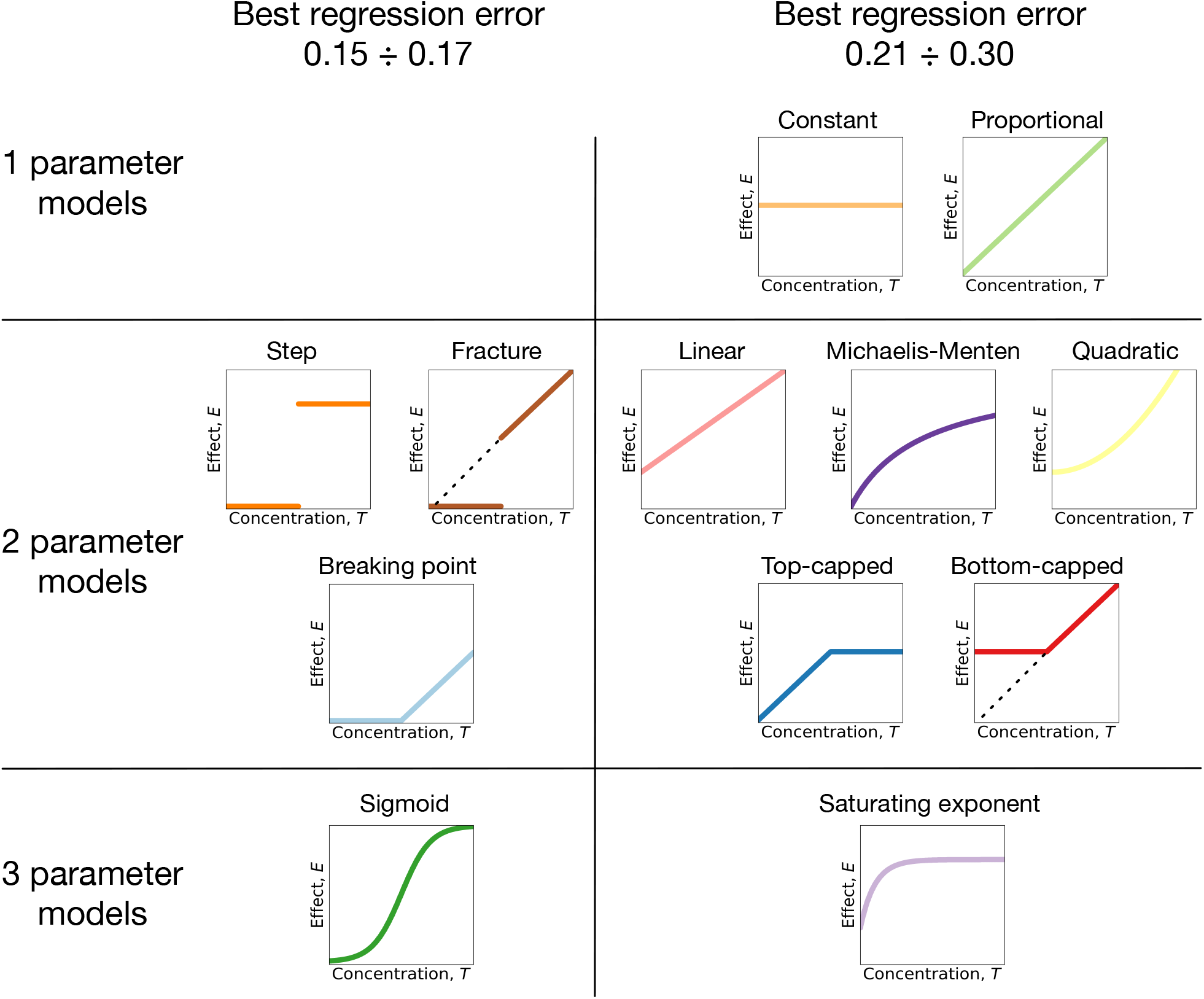
Models of the acting substance concentration effect in the disconnecting and toxic compound models. We consider twelve models of the relationship between acting substance concentration and its effect on the filaments. Two models have a single parameter each and serve as a control. Eight models have two parameters. Two remaining models have three parameters each, see Table S2 for details. Regression errors are shown for the disconnecting compound models. There, four models: step, fracture, breaking point, and sigmoid have shown much smaller regression errors than other models.

**Figure S5.**
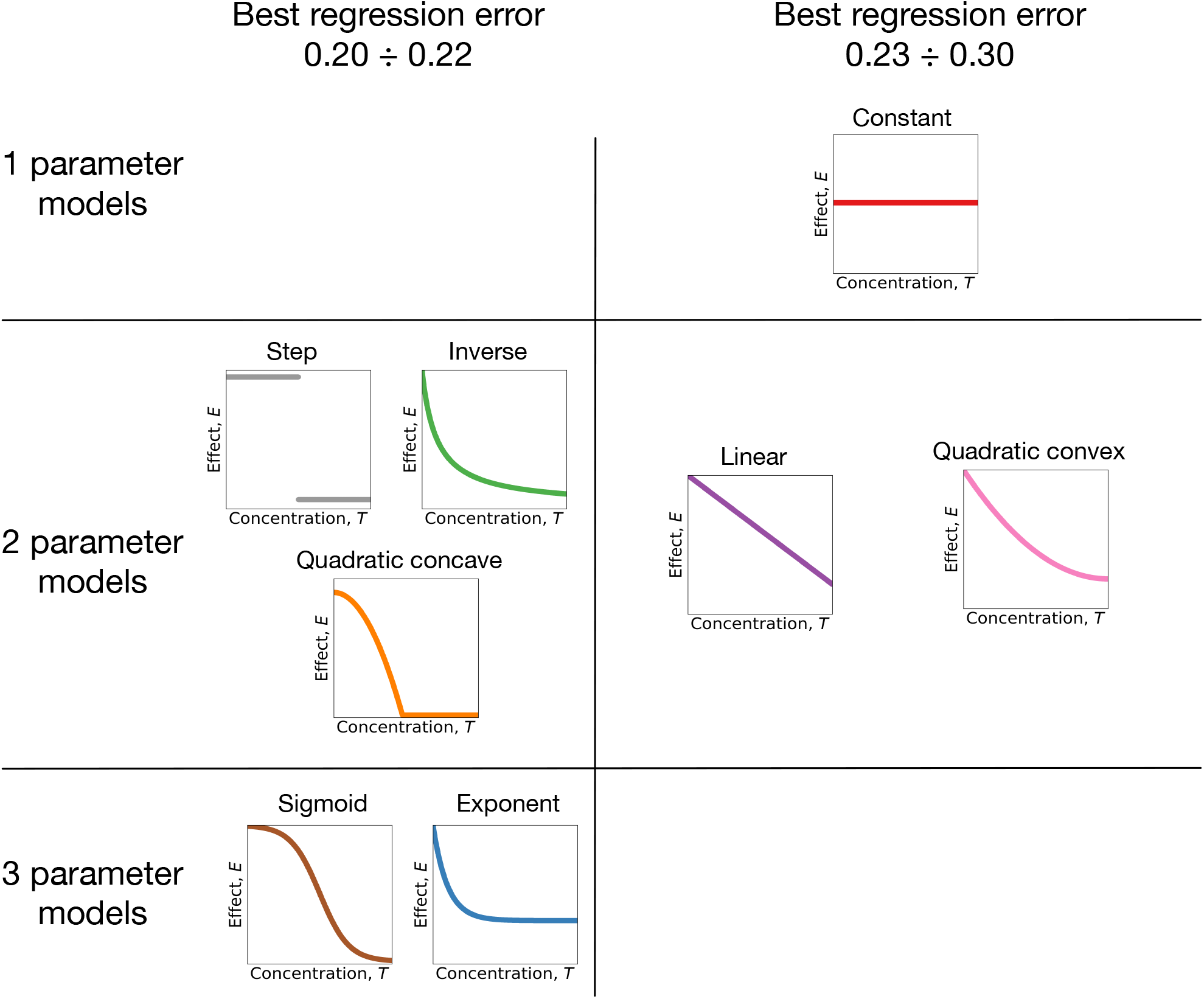
Models of the acting substance concentration effect in the connecting compound models. We consider eight models of the relationship between acting substance concentration and its effect on the filaments. One model has a single parameter and serves as a control. Five models have two parameters. Two remaining models have three parameters each, see Table S3 for details. There, five models: step, inverse, quadratic concave, exponent, and sigmoid have similar small regression errors.

**Table S2.**
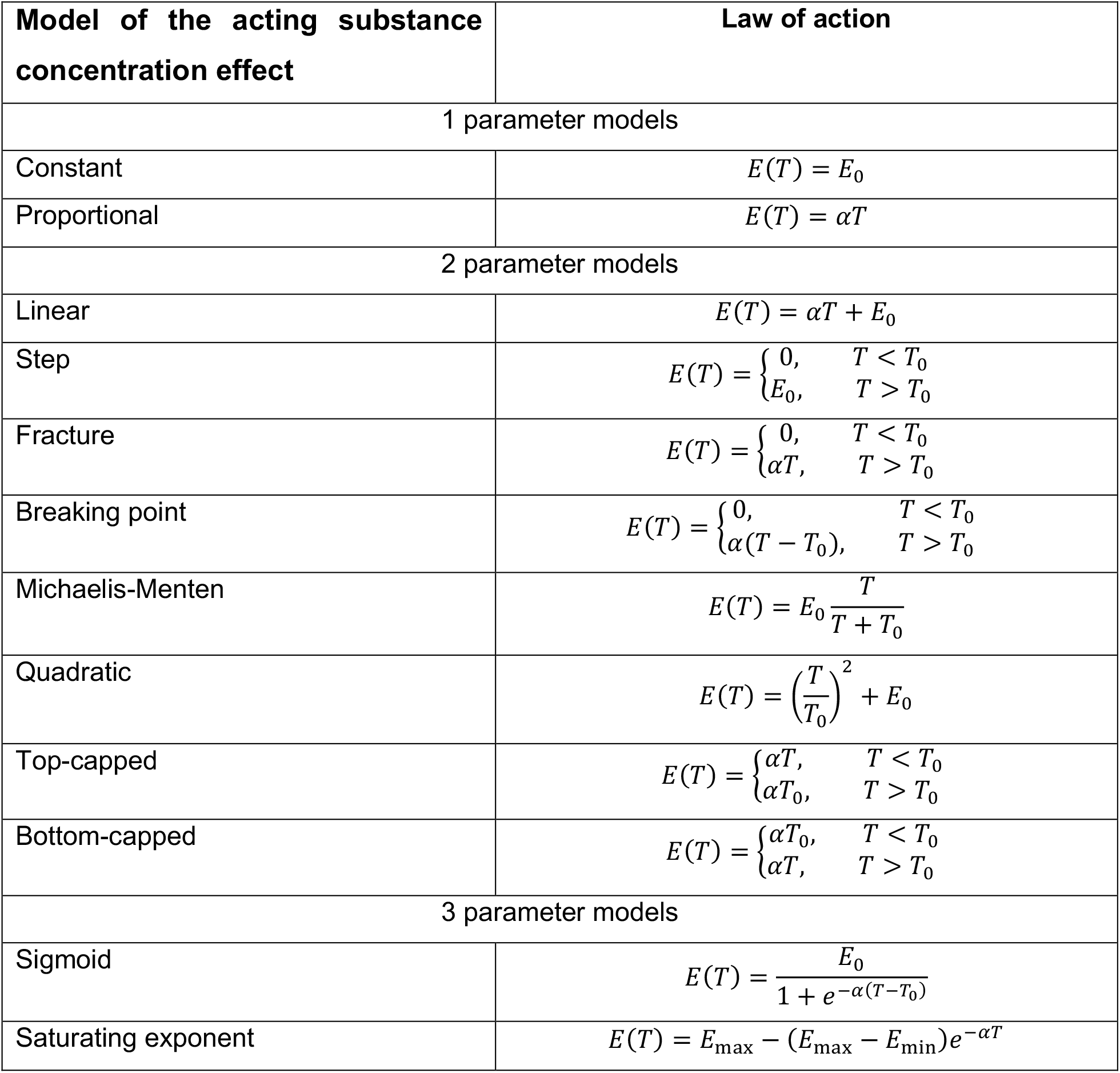
Action law E(T) in models used in the toxic and disconnecting compound model families.

**Table S3.** Action law E(T) in models used in the connecting compound models family.

| Model of the acting substance<br>concentration effect | Law of action |
| --- | --- |
| 1 parameter models |  |
| Constant | $E(T) = E_0$ |
| 2 parameter models |  |
| Linear | $E(T) = \alpha(1 - T) + E_0$ |
| Step | $E(T) = \begin{cases} E_0, & T < T_0 \\ 0, & T > T_0 \end{cases}$ |
| Quadratic convex | $E(T) = E_0 + \alpha(1 - T)^2$ |
| Quadratic concave | $E(T) = \max(0, E_0 - \alpha T^2)$ |
| Inverse | $E(T) = \frac{E_0}{1 + \frac{T}{T_0}}$ |
| 3 parameter models |  |
| Sigmoid | $E(T) = \frac{E_0}{1 + e^{\alpha(T-T_0)}}$ |
| Decaying exponent | $E(T) = E_{\max} + (E_{\max} - E_{\min})e^{-\alpha T}$ |

**Figure S6.**
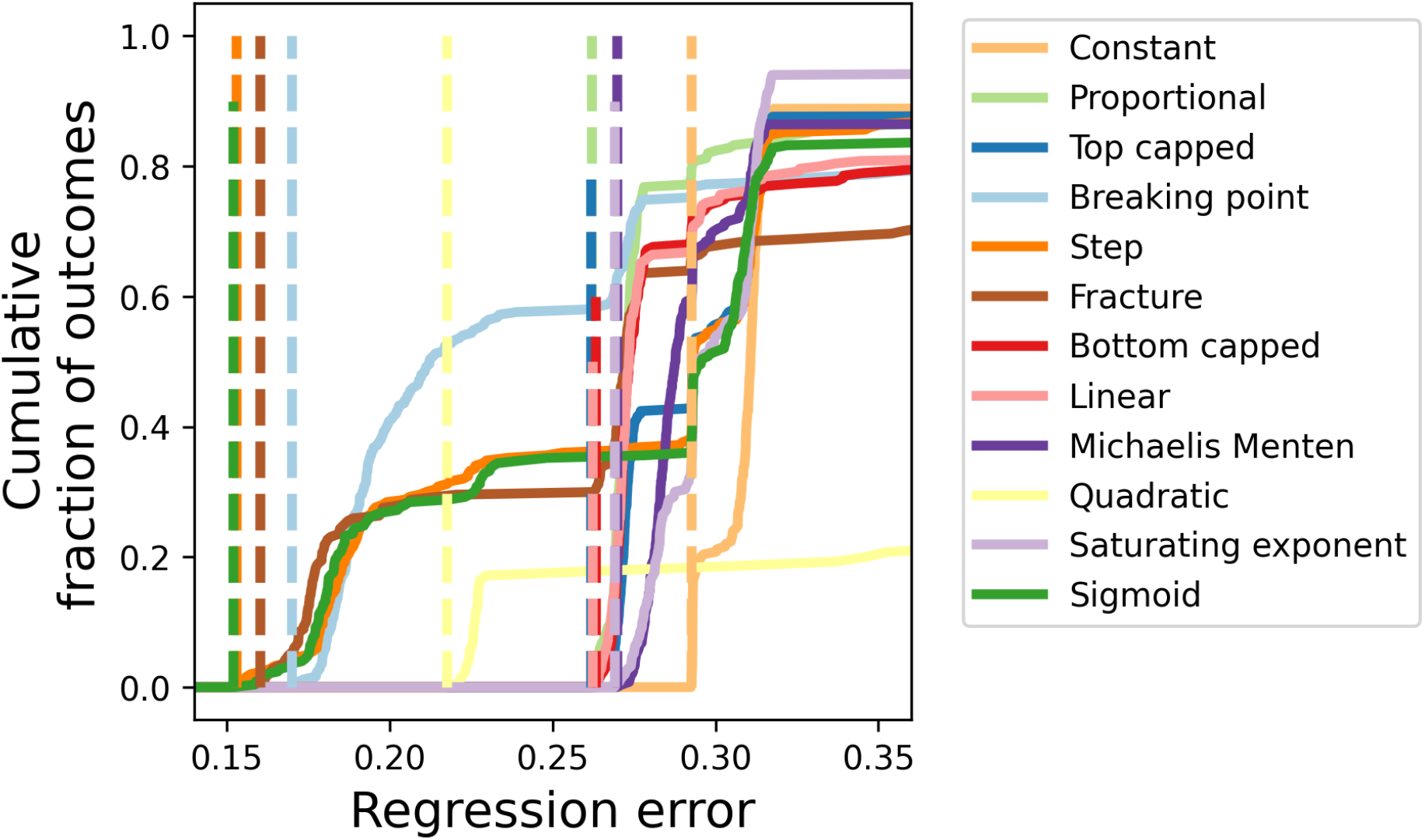
Cumulative distribution functions and the minimal regression errors obtained for each of twelve disconnecting compound models. Studied models can be classified into two groups: models with a good fit having minimal regression errors below 0.17, and models with worse fit, for which the minimal regression error is above 0.21 (can be increased to 0.26 if quadratic model is dropped), see also Table S4. Plots show sample cumulative distribution functions of regression errors from 250 independent optimizations for each model. Dashed lines represent the minimal regression error in each model.

**Figure S7.**
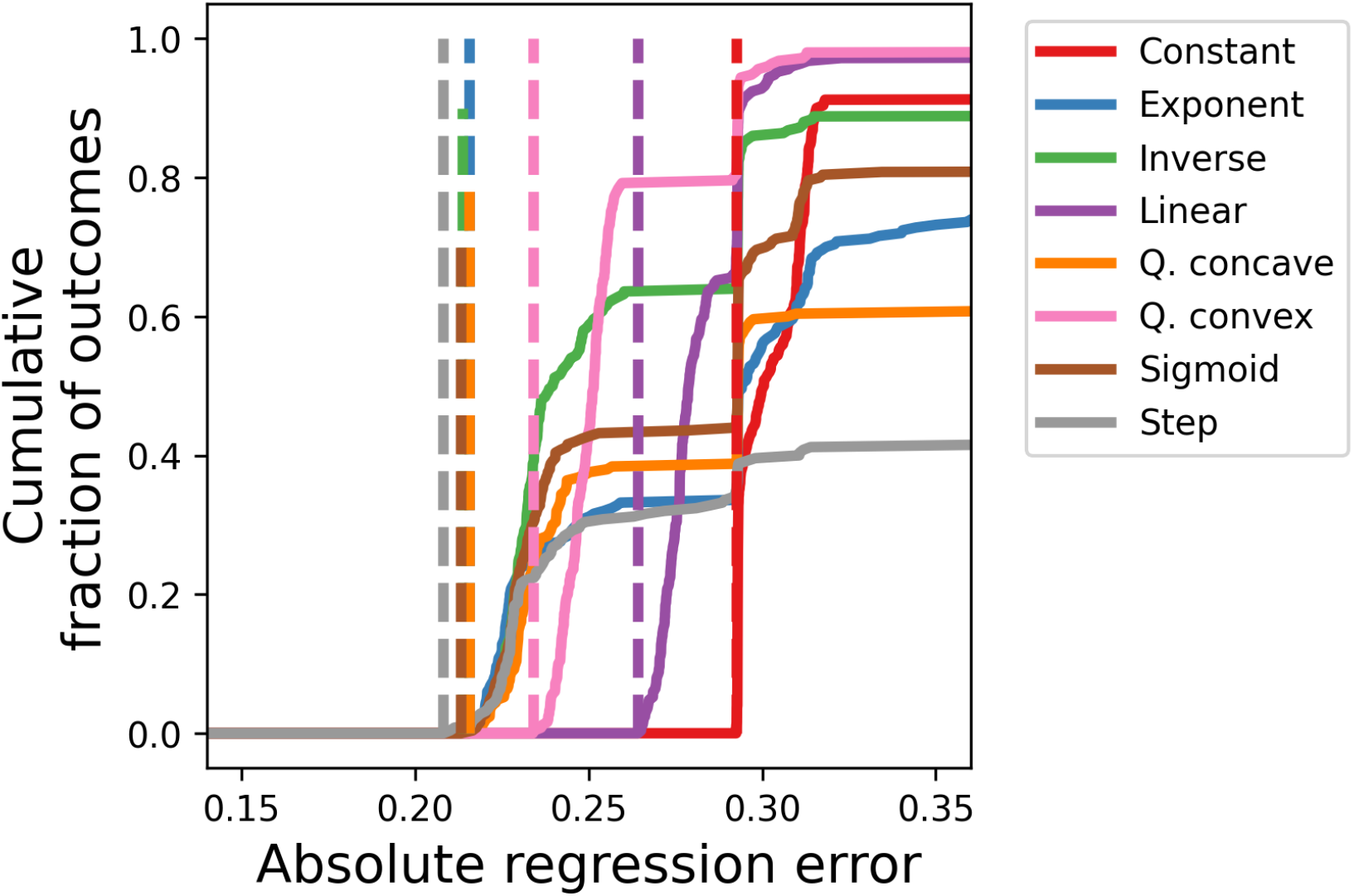
Cumulative distribution functions and the minimal regression errors obtained for each of the eight connecting compound models. Studied models can be classified into two groups: models with a good fit having minimal regression errors around 0.21, and models with worse fits, for which the minimal regression error is above 0.22, see also Table S5. Plots show sample cumulative distribution functions of regression errors from 250 independent optimizations for each model. Dashed lines represent the minimal regression error in each model.

**Table S4.** Minimal regression errors obtained for the disconnecting and toxic compound models across 250 independent optimizations. Models are sorted by the minimal regression error in the disconnecting compound family. Models with the highest quality fitting are highlighted.

| Models | Disconnecting compound<br>regression error | Toxic compound<br>regression error |
| --- | --- | --- |
| Sigmoid | 0.152 | 0.583 |
| Step | 0.153 | 0.594 |
| Fracture | 0.160 | 0.721 |
| Breaking point | 0.170 | 0.724 |
| Quadratic | 0.217 | 0.583 |
| Proportional | 0.262 | 0.732 |
| Linear | 0.262 | 0.596 |
| Top-capped | 0.262 | 0.593 |
| Bottom-capped | 0.263 | 0.613 |
| Michaelis-Menten | 0.270 | 0.598 |
| Saturating exponent | 0.269 | 0.560 |
| Constant | 0.292 | 0.590 |

**Table S5.** Minimal regression errors obtained for connecting compound models across 250 independent optimizations. Models with the highest quality fitting are highlighted.

| Models | Connecting compound<br>regression error |
| --- | --- |
| Step | 0.208 |
| Sigmoid | 0.213 |
| Inverse | 0.214 |
| Quadratic concave | 0.216 |
| Exponent | 0.216 |
| Quadratic convex | 0.234 |
| Linear | 0.264 |
| Constant | 0.292 |

### Text S1. Comparison of our study to the cyanobacterial filament fragmentation model by Rossetti et al (2011)

A model of filament fragmentation is also described in Rossetti et al (2011). Their model can be considered as a specific limited case of our set of models. Both our and Rossetti et al models examine growing populations of linear filaments and share the same logistic dependence of the cell division rate from the cell density. However, the model of Rossetti et al includes only filament fragmentation due to cell death, while our model set also includes the mechanism of connection loss.

Another difference between the models is that in our case, cell death is caused by the compound produced by cells; while in the model of Rossetti et al the rate of cell death depends on cell density. In both cases, more cells in a population results in a larger death rate but the model presented here features a reactivity: a sudden increase in cell count does not cause an immediate increase in death rate; instead the death rate will steadily rise with the accumulation of the compound. This difference is of principal importance: the model of Rossetti et al is unable to recover the results of supernatant experiments presented in Fig. 2 – the media plays no role in that model and it is impossible to observe high rates of filaments fragmentation at low cell densities. Nevertheless, the design of the cell death from the model of Rossetti et al can be formally recovered by our models set by choosing the proportional model of compound action (see Table S2 and Fig. S4), plus setting both compound decay rate *D*_comp_, and compound toxicity *α* to very large values. With a high compound decay rate, the equilibration of the compound concentration will occur rapidly, so the death rate will closely follow the cell density. Since a high decay rate also means low overall concentrations of compound, to have any significance, the toxicity must also be high. With this set up our model will behave identically to the model of Rossetti et al, however by doing so, the choice of the compound action mechanism, the model of the compound action, and even the parameter values would be far from optimal.

### Text S2. Half of the stationary population, where filaments disintegrate due to cell death must be filamentous

The toxic compound models assume that a filament breaks because a cell in it dies and can no longer hold the two side branches together. These events occur more frequently at higher concentrations of the compound – so the filaments that make up the population become shorter as the compound concentration increases in the environment. However, this process cannot bring a population to a state where only single cells exist – the observation we have made in all our experiments. This is because the concentration of the toxic compound cannot be arbitrarily high – too much compound would lead to a population decline. The maximum possible concentration of the compound (and the shortest possible filaments) is reached at the equilibrium between growth and death, when the growth of the population due to cell division is compensated for by the loss of cells through the action of the toxic compound.

Consider this equilibrium in detail, with a particular focus on solitary cells. Solitary cells arise in the population through the fragmentation of filaments after the death of a cell. There are also two processes by which solitary cells are removed from the population: Cell death due to exposure to the toxic compound and cell division, which turns a single cell into a two-celled filament. In the stationary state, the rates of gain and loss of solitary cells are equal

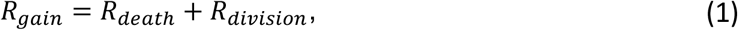

where *R*_*gai*n_ is the rate of gain of solitary cells, *R*_*death*_ is the total death rate of solitary cells, and *R*_*divi*s*i*on_ is the total division rate of all solitary cells (all rates are per population, not per cell). Next, we elaborate on these rates. Let us denote the number of solitary cells in a population as *N*_1_ and the number of filaments as *N*_*F*_. If the division rate for a single cell is *r*, then the total division rate for all solitary cells is

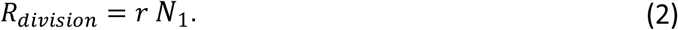

The stationary state of the population is defined by the equilibrium between cell growth (both solitary cells and filaments) and cell death, i.e. for each cell the rates of death and division are equal. Therefore, in this state, the rate of cell death is also *r*, so that the total rate of death among the solitary cells is

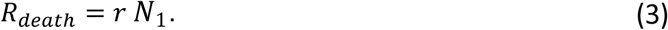

Finally, the gain of solitary cells is determined by the fragmentation of filaments. There might be filaments of various lengths in the population. However, for a filament of any length, there are only two cells, which deaths release a solitary cell, i.e. the second cell at each tip. Therefore, the rate of gain is

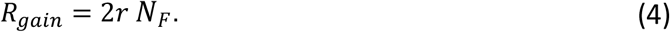

Combining Eqs. 1-4, we get

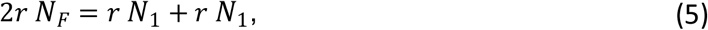

From here, it immediately follows that

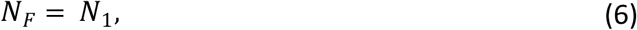

i.e. the number of filaments in the stationary state is equal to the number of solitary cells. This proves that the fragmentation of the filaments caused by the toxic compound cannot lead to a stable population consisting of only solitary cells, as we observe in our experiments.

**Figure S8.**
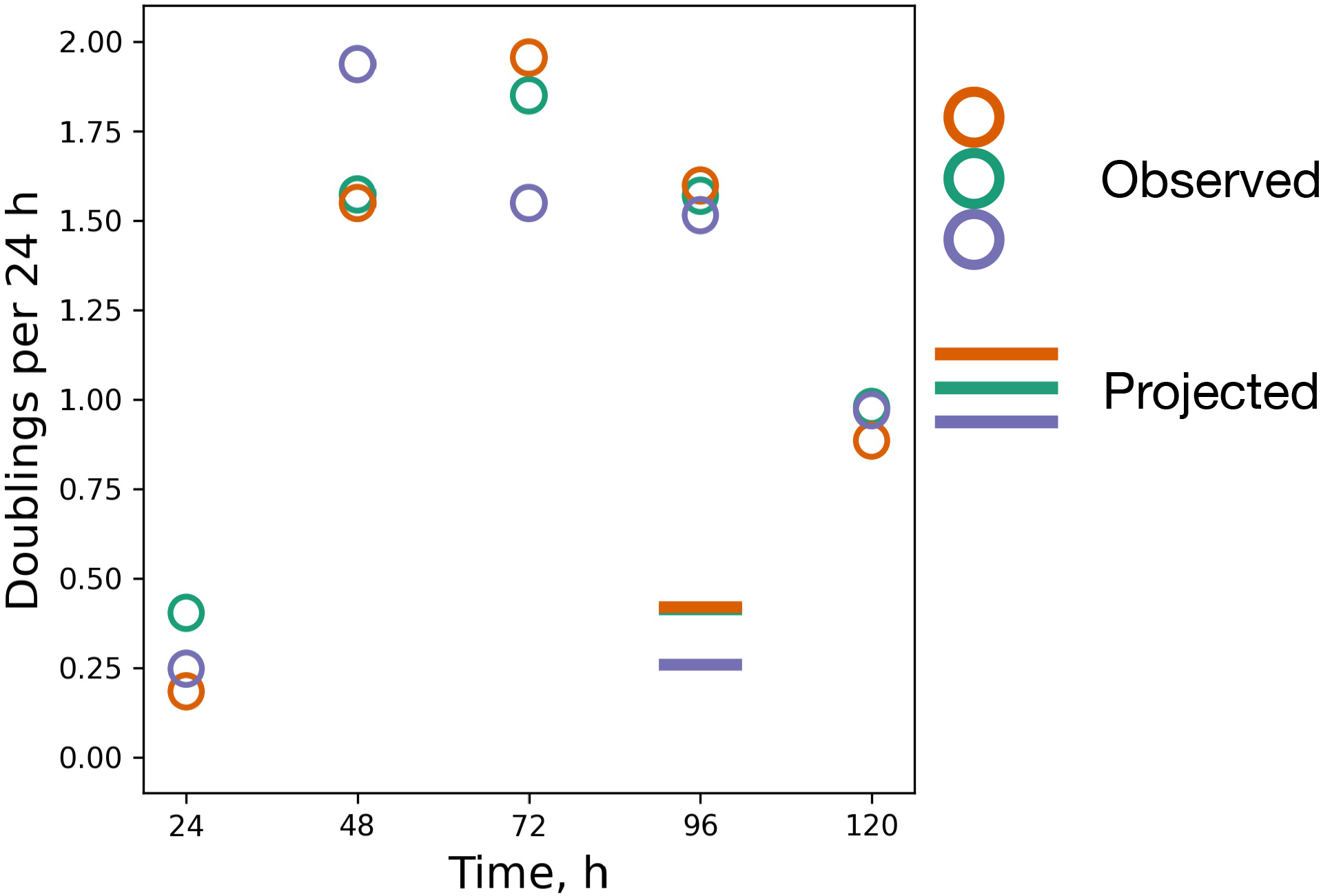
The population growth rate does not change at the time the filaments dissolve. The figure shows the growth rates of the cyanobacterial populations as a function of time. The colored circles are data from experimental replicates (the value plotted at X hours is the number of cell doublings between X-24 and X hours). The colored lines at 96 h (the timepoint where the filaments have dissolved into solitary cells) show what the population growth rate would be if every second cell in a filament died and the remaining cells were released as single cells. These predicted values are much lower than those actually observed. The fragmentation was therefore not caused by a mass death of cells.

**Figure S9.**
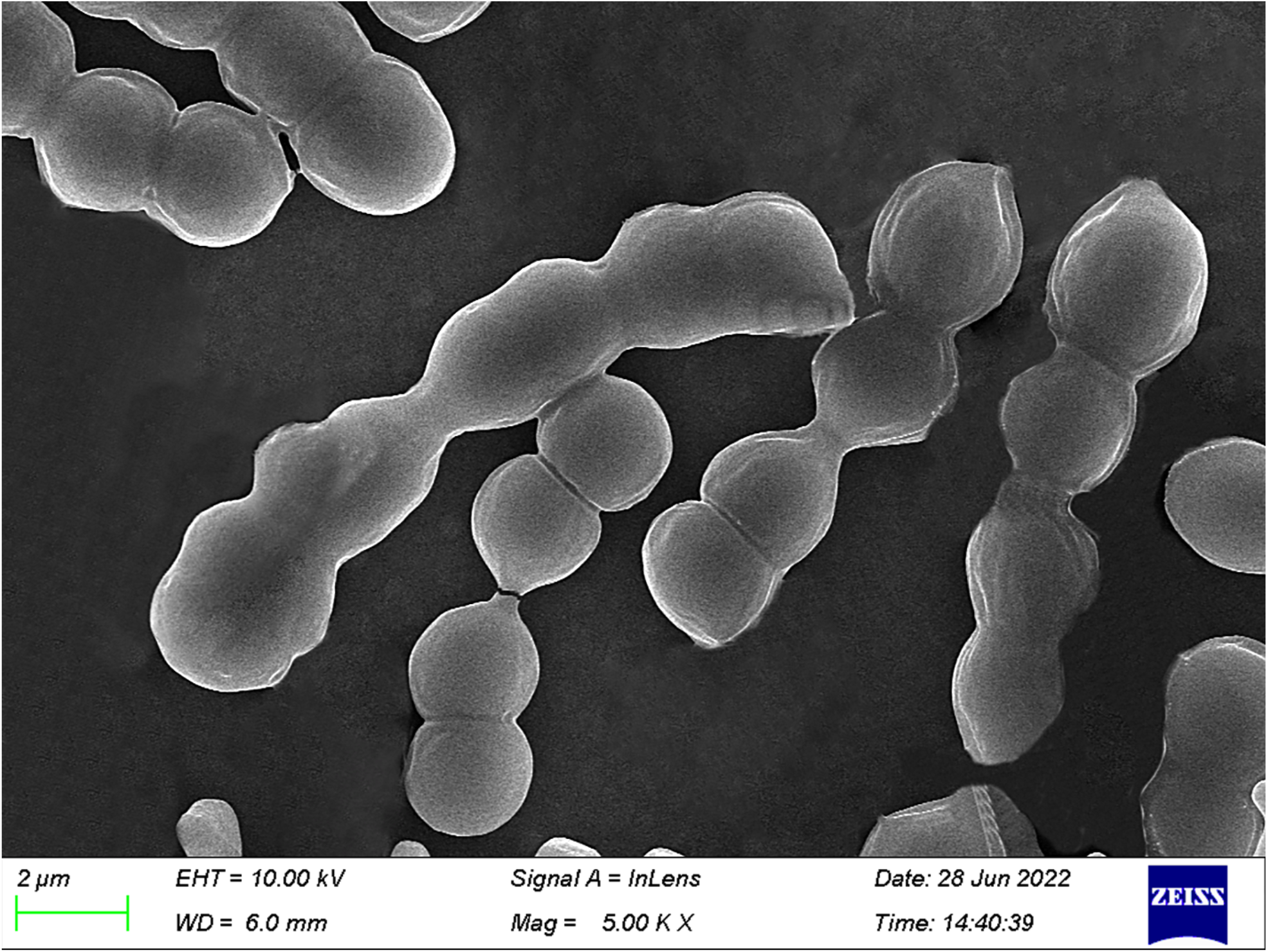
Scanning electron photomicrographs of 72-hour-old filaments of *Cyanothece* sp. ATCC 51142. Note the breaking cell-cell connection in the center filament.

